# Perspective/Review: Regulating telomere length from the inside out: The replication fork model

**DOI:** 10.1101/041772

**Authors:** Carol W. Greider

## Abstract

Telomere length is regulated around an equilibrium set point. Telomeres shorten during replication and are lengthened by telomerase. Disruption of the length equilibrium leads to disease, thus it is important to understand the mechanisms that regulate length at the molecular level. The prevailing protein counting model for regulating telomerase access to elongate the telomere does not explain accumulating evidence of a role of DNA replication in telomere length regulation. Here I present an alternative model: the replication fork model that can explain how passage of a replication fork and regulation of origin firing affect telomere length.

## Introduction

Telomere length homeostasis is essential for cell survival. Short telomeres trigger DNA damage, induce cellular senescence and apoptosis, and cause Short Telomere Syndromes and age-related disease (Armanios, 2009). Cancer cells, on the other hand, maintain or elongate telomeres and escape senescence to allow immortal growth (Greider, 1999). Telomeres naturally shorten during DNA replication, which is counterbalanced by *de novo* addition of telomere sequences by telomerase (Greider and Blackburn, 1985). Most of the telomere is replicated by conventional replication machinery (Wellinger and Zakian, 2012); however, at each cell cycle telomerase elongates a few telomeres by addition of a few repeats (Teixeira et al., 2004). The central question is, what determines whether a telomere will be elongated and how does this establish length homeostasis? Here I present a model for how the stochastic elongation of a few telomeres at each cell cycle can be explained by coupling between DNA replication and telomere length maintenance.

### Telomere binding proteins regulate telomere length

Telomeres are made up of simple G-rich DNA sequence repeats that are packaged into chromatin (Tommerup et al., 1994) and bound by telomere specific binding proteins. In mammalian cells, the shelterin complex consists of TRF1 and TRF2, which bind along the double stranded telomere sequence and recruit associated proteins TIN2, TPP1, POT1 and RAP1 (Palm and de Lange, 2008). POT1 binds tightly to the single stranded G rich telomere DNA sequence. Telomeres in *S. cerevisiae* were initially reported to be non-nucleosomal (Wright et al., 1992), however recent data suggests nucleosomal packaging in yeast as well (Pisano et al., 2008; Rossetti et al., 2001). In *S. cerevisiae*, the Rap1 protein binds to the double stranded telomere repeats and either Rif1 and Rif2, or Sir3 and Sir4, bind to the C-terminal domain of Rap1 (Shore and Bianchi, 2009). The single-stranded G-rich telomeric DNA is bound by Cdc13, (Lin and Zakian, 1996; Nugent et al., 1996) and the associated Stn1 and Ten1 proteins (Grandin et al., 2001; Grandin et al., 1997). The double stranded and single stranded telomere specific binding proteins are essential for *both* the protection of the chromosome end and for regulating telomerase access to the telomere (Palm and de Lange, 2008; Wellinger and Zakian, 2012). How they carry out these functions is critical to understanding length regulation.

### Protein counting model

Two experimental findings helped establish the “protein-counting model” for telomere length regulation (Marcand et al., 1997). First, in *S. cerevisiae*, both C-terminal mutations in *RAP1* (Sussel and Shore, 1991), or deletion of the genes encoding two Rap1 interacting proteins, *RIF1* and *RIF2*, cause excessive telomere elongation (Hardy et al., 1992; Wotton and Shore, 1997) implying these proteins normally block telomere elongation. Second, short telomeres are more likely to be elongated by telomerase than long telomeres (Marcand et al., 1999; Teixeira et al., 2004). To account for these results and others, the ‘protein counting’ model for telomere length regulation (Figure 1A) was proposed in 1997 (Marcand et al., 1997). This model was also adapted to explain human telomere length regulation, since knockdown of the telomere binding proteins TRF1, TRF2, POT1, and TIN2 also caused excessive telomere elongation (Loayza and De Lange, 2003; Takai et al., 2010; van Steensel and de Lange, 1997; Ye and De Lange, 2004). The evolutionarily conservation of negative telomere length regulation by telomere binding proteins helped solidify the protein counting model (Smogorzewska et al., 2000).

**Figure 1.**
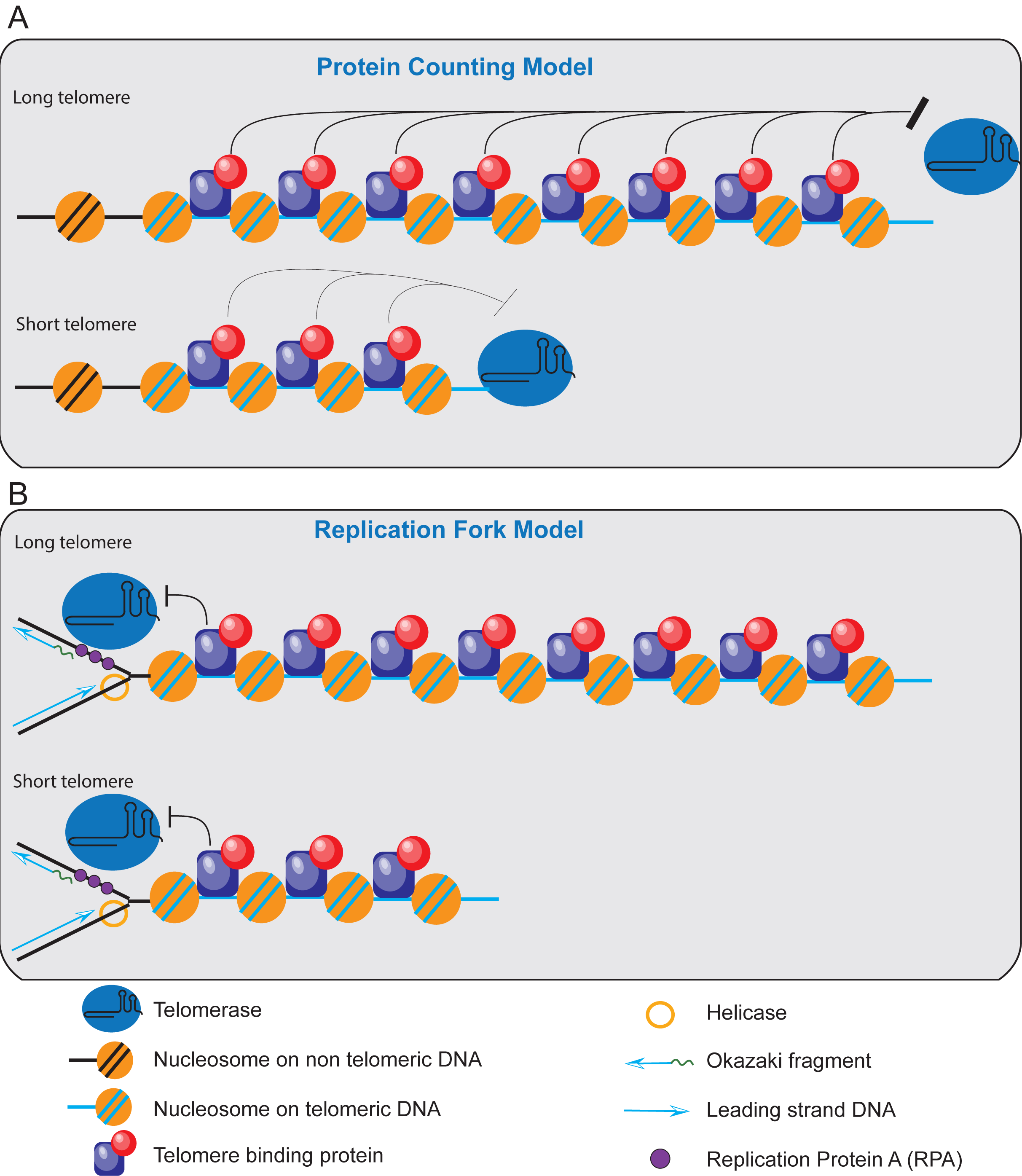
Old and new models for regulating elongation of telomeres by telomerase. *[A]* Counting model for telomere length regulation. Telomeric DNA (light blue line) is shown packaged as nucleosomes (orange circles) and bound by interspersed telomere specific proteins (blue and red). The telomere proteins act from a distance to block telomerase access to the end of the chromosome. The long telomere (top) has greater repressive effects (black bar) on telomerase (green oval) than the short telomere. *[B]* Replication fork model for telomere elongation. Telomerase is shown traveling with the lagging strand machinery. The RPA or t-RPA is shown in purple, and the helicase is shown as a yellow ring. The fork replicates through nucleosomes and bound telomere proteins, both of which can cause dissociation of telomerase from the fork. Telomerase must remain bound to the fork as it reaches the extreme terminus for the telomere to be extended.

At its core, the protein counting model states that there is an additive negative effect of telomere bound proteins on telomerase access to the telomere. That is, long telomeres have a stronger repressive effect that keeps telomerase off the 3’ end of the telomere, while short telomeres have a weaker repressive effect and so telomerase can elongate them (Figure 1A). Although this model explains the negative inhibitory role of telomere binding proteins, it is unclear mechanistically how an additive negative effect might be integrated and/or propagated along many kilobases of telomere sequence. Additionally it is unclear how the accumulated bound proteins block telomerase access to the very terminus. This protein-counting model does also not explain a number of new experimental findings, as discussed below, suggesting that alternative models should be considered.

### A Replication fork model for telomere length regulation

An alternative model for telomere length regulation better accounts for new (and old) research linking DNA replication and telomere elongation. This ‘replication fork model’ accounts for both negative regulation of telomere elongation and preferential elongation of short telomeres. In this model, telomerase is recruited to the end of the telomere through an association with the replication fork (Figure 1B). Telomere-binding proteins exert their negative effect by increasing the probability that telomerase will dissociate from the traveling replication fork. Therefore the longer the telomere, the lower the probability of telomerase reaching the end where it can preform its catalytic function. As the fork progresses toward the telomere, each bound protein that the fork encounters results in a small probability of telomerase dissociating from the replication fork. The nucleosomes that the fork encounters might exert some negative effect and, in addition, this model proposes that replicating past telomere specific binding proteins increase the probability that telomerase will dissociate from the fork. On a longer telomere the cumulative small probabilities of telomerase dissociation make it less likely telomerase will arrive at the terminus. This model fits the long established evidence that short telomeres are preferentially elongated and that telomere elongation is stochastic; only a few telomeres are elongated at every cell cycle. This model also explains how telomere-binding proteins negatively regulate telomere length, they may provide a simple barrier, like the nucleosome, or some may actively promote dissociation of telomerase from the fork.

There are a precedents for proteins traveling with the replication fork, including Mrc1 and Tof1 that form the <u>f</u>ork <u>p</u>rogression <u>c</u>omplex (Katou et al., 2003). DDK travels with the replication fork to regulate double strand breaks in meiosis (Murakami and Keeney, 2014), and RRM3 travels with the fork to promote replication thought specific barriers (Azvolinsky et al., 2006). The FACT complex involved in chromatin remodeling, and Dia2 involved in replication termination are also tethered to the replisome (Foltman et al., 2013; Morohashi et al., 2009).

There is early evidence from ciliates that telomerase also travels with the replication fork. In hypotrichous ciliates, replication initiation and progression is coordinated across the macronucleus in a ‘replication band’ (Olins et al., 1989). This band progresses synchronously across the nucleus synthesizing DNA. The Cech labs showed that telomerase associates with these replication bands in *Oxytricha* as it travels with the replication forks during S phase (Fang and Cech, 1995). The coordination of replication fork progression and telomerase delivery at to the very end would help explain why telomerase elongates telomeres at the very end of S phase.

The relative stoichiometries of telomerase and replication forks may explain the stochastic nature of telomere elongation. The concentration of telomerase *in vivo* is very low; in *S. cerevisiae* there are about 20 copies and in human cancer cell lines about 250 copies of telomerase per cell (Mozdy and Cech, 2006; Xi and Cech, 2014). In *S. cerevisiae* there are approximately 626 unique origins as well over 200 rDNA origins, a subset of which fire each cell cycle (Siow et al., 2012), thus telomerase might only associate with a small fraction of the forks as they travel to the ends of chromosomes. If telomerase has some probability of associating with all replication forks, this could explain telomere repeat addition at internal break sites. This form of “chromosome healing” occurs with some frequency in many organisms including humans and *S. cerevisiae* (Greider, 1991; Myung et al., 2001; Ribeyre and Shore, 2013). As discussed below, telomerase may also have a higher probability of associating with telomeric forks since it binds to an alternative, telomere specific, RPA.

### Origin placement and firing efficiency could regulate telomere length

The replication fork model provides a plausible explanation of two previously mystifying results: how subtelomeric sequences and the regulation of origin firing both affect telomere length. Both origin location, and non-firing of a telomeric origin, will affect how far a fork must travel before it reaches the chromosome end (Figure 2). The probability that telomerase will remain bound to the replication fork until the end of the chromosome will increase with a shorter distance between the most telomere-proximal origin and the chromosome end.

**Figure 2.**
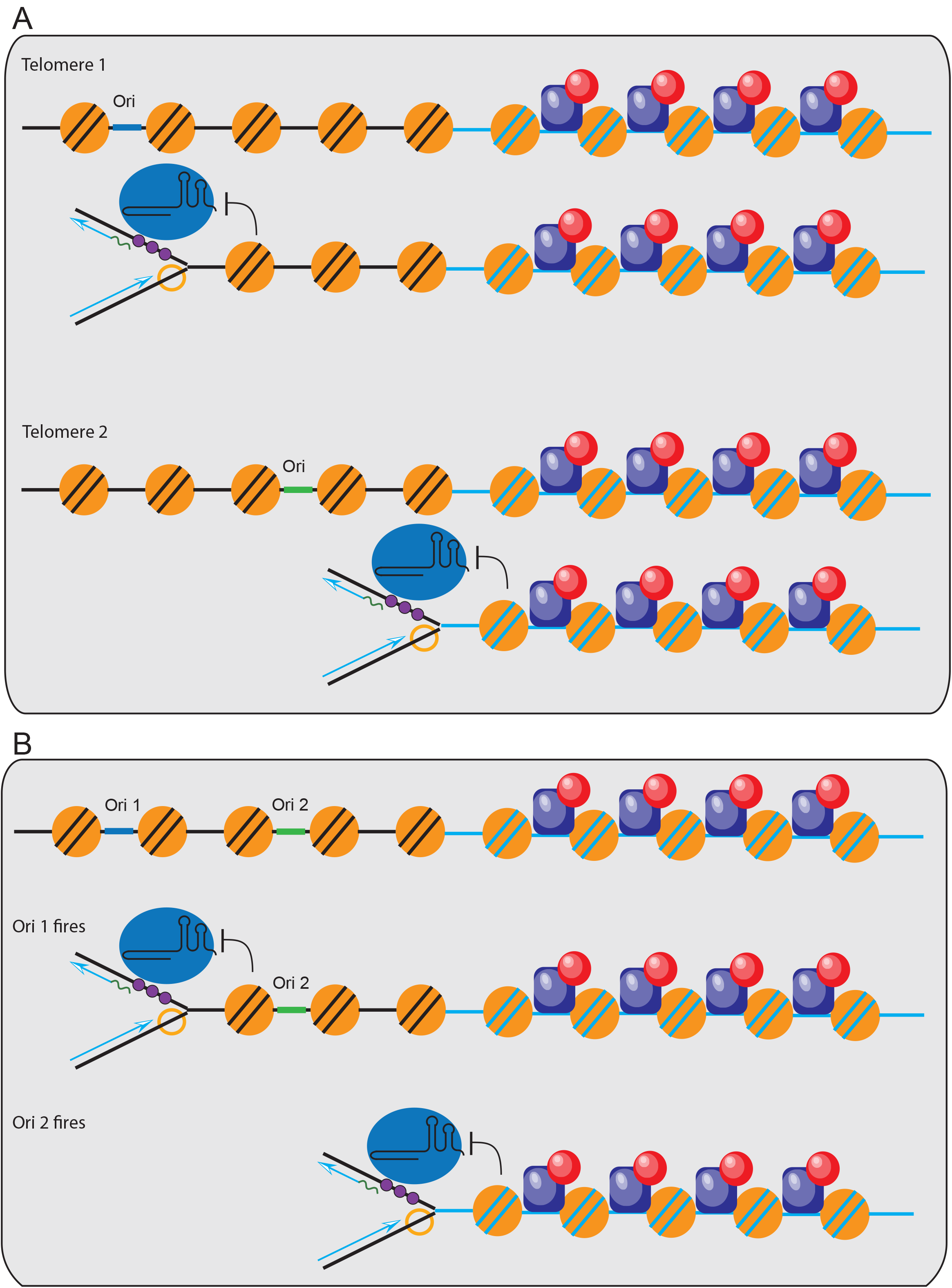
Distance from an origin may affect telomere length. *[A]* If telomerase associated with the origin on telomere 1 that is more distant from the end of the chromosome, there is a high probability of dissociating before reaching the chromosome end. In contrast, on telomere 2 the origin is more proximal to the end of and thus this telomere has a higher probability of being elongated. *[B]* Telomere proximal origins are inhibited from firing and can be passively replicated by adjacent origins. Here Ori 1 is efficient while Ori 2 does not fire in every cell cycle. If travels with the fork that initiates at Ori1, the probability of it reaching the end is relatively low. In contrast, if Ori 2 fires there is a shorter distance to the chromosome end, and telomerase has a high probability of elongating that telomere. Rif1 normally blocks the telomeric Ori 2 from firing; in the absence of Rif1, Ori 2 will fire and telomeres elongate.

The differential locations of origins may explain a curious discovery in the Petes lab that the length of telomeres containing Y’ subtelomeric sequence are regulated differently than telomeres containing subtelomeric X sequence (Craven and Petes, 1999). Telomeres in *S. cerevisiae* contain two types of repetitive subtelomeric sequences, termed X or Y’, immediately adjacent to the G rich telomere repeats (Chan and Tye, 1983). Both of these elements contain replication origins, but the distance of the origin from the chromosome end varies in the two repeats (Louis, 1995). The replication fork model of telomere length regulation would suggest that differential proximity to an origin in X and Y’ containing telomeres could result in different probabilities of telomere elongation (Figure 2A).

### Replication origin firing regulates telomere length

A second curious finding that can be explained by the replication fork model is the role of Rif1 in regulating origin firing and telomere length. Telomeric origins replicate late in S phase and often do not file and are passively replicated (McCarroll and Fangman, 1988; Raghuraman et al., 2001). Strikingly, it is the telomeric location, not the DNA sequence of the origins, that determines their firing efficacy (Ferguson and Fangman, 1992). If an early firing origin from elsewhere in the genome is relocated to the telomere, it will now fire late, or not at all. Conversely, a telomeric origin placed on a circular plasmid will fire early and efficiently. Strikingly this late replication of a telomeric origin is conserved in human cells (Smith and Higgs, 1999). New results directly link Rif1 to this regulation of telomeric origin firing.

Rif1 was first identified in *S. cerevisiae* as a negative telomere length regulator, and helped form the basis for the protein-counting model (Hardy et al., 1992; Levy and Blackburn, 2004; Marcand et al., 1997). Experiments from several different groups now show that Rif1 is an evolutionarily conserved regulator of origin firing. Deletion of *RIF1* in yeast, or knockdown in mammalian cells, allows the origins that were blocked in early S phase to now fire (Buonomo et al., 2009; Cornacchia et al., 2012; Hayano et al., 2012; Lian et al., 2011; Mattarocci et al., 2014; Peace et al., 2014; Sreesankar et al., 2015; Yamazaki et al., 2012). Rif1 blocks the origin firing through recruitment of Protein Phosphatase 1 (PP1), which in turn antagonizes the action of the DDK1 kinase, required for origin firing (Dave et al., 2014; Hiraga et al., 2014; Mattarocci et al., 2014). Rif1 bound at telomere repeats will thus recruit PP1 and inhibit firing of origins near the telomere. Longer telomeres would have more bound Rif1 leading to increased PP1 recruitment and decreased firing of telomere proximal origins and late origin firing will be inhibited near telomeres.

The replication model suggests how increasing the probability of telomeric origin firing can lead to longer telomeres in a *RIF1* deletion mutant. If the telomere proximal origin does not fire due to Rif1/PP1 activity, then replication origins will have to come from the next most internal origin (Figure 2B). However when *RIF1* is deleted the telomere proximal origins fire, thus decreasing the distance to the chromosome end and increasing the probability that telomerase will elongate the telomere. The replication fork model thus links the long telomeres in *RIF1* deletion mutants with its effect on origin firing.

### Feedback loop for origin activation and repression may regulate telomere length

Telomere length homeostasis may be established by a feedback loop between origin firing efficacy and telomere length (Figure 3). The increased recruitment of PP1 to long telomeres decreases the probability of adjacent origin firing, and thus over many cell cycles long telomeres will shorten due to the end replication problem. When that telomere becomes shorter, there will be less Rif1 bound and the telomere proximal origin can fire again, thus increasing the probability that telomerase will arrive at the end to extend that telomere. Interestingly, when a telomere is artificially shortened the telomere-proximal origin fires more efficiently (Bianchi and Shore, 2007a), supporting a feedback mechanism between telomere length, origin activity and telomere elongation (Figure 3). Having outlined the general concept of the replication fork model of telomere elongation, I will now examine how previous experiments that connected replication and telomere length can be interpreted in light of this model.

**Figure 3.**
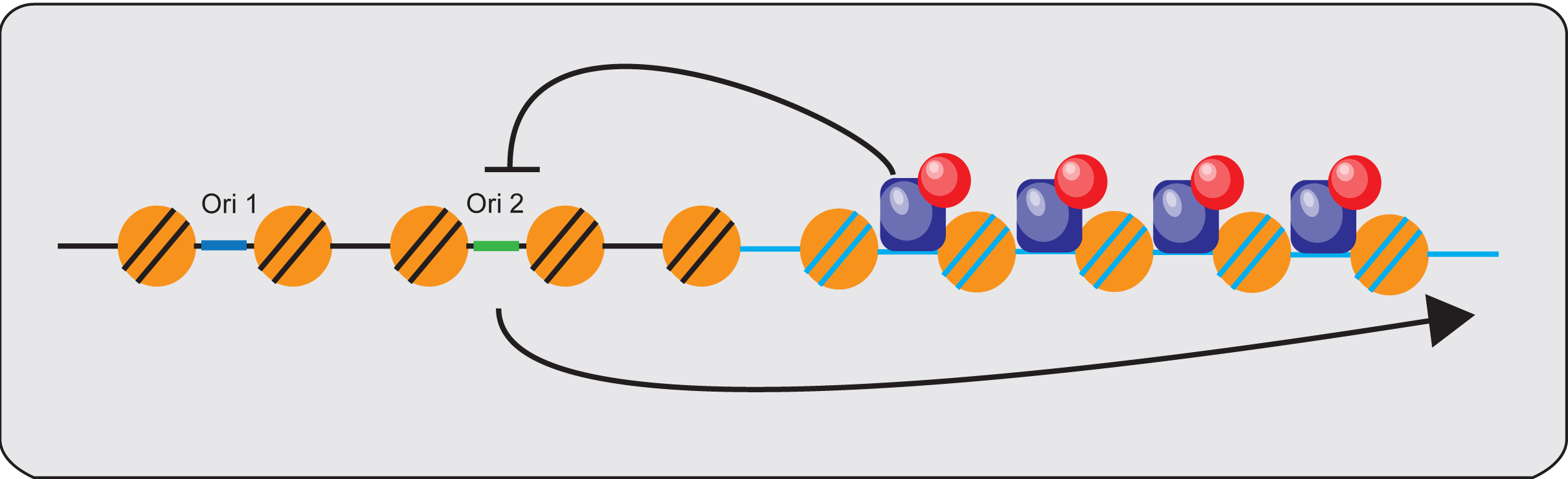
Feedback regulation of origin firing maintains telomere length homeostasis. At long telomeres, local Rif1 binding to telomere DNA (blue lines) blocks origin firing at proximal telomeres in adjacent DNA (black lines). The telomere is then replicated from the more distal Ori 1. At short telomeres the fewer binding sites for Rif1 allows Ori 2 firing and this increases the probability of telomere extension.

### Telomere elongation is linked to replication

The association of telomere length changes with DNA replication has been noted for some time and, in fact, some elements of this model have been previously suggested in the literature. Wellinger proposed that origin firing is coupled to telomere elongation (Wellinger et al., 1993) and also that telomere elongation requires passage of a replication fork (Dionne and Wellinger, 1998). Other groups have also linked origin firing to telomere elongation; by following elongation of an artificially shortened telomere in *S. cerevisiae* through the cell cycle, two groups found that telomere elongation coincides exactly with telomere replication (Bianchi and Shore, 2007a; Marcand et al., 2000). In human cells, following replication of a single telomere showed telomerase elongation occurs immediately following replication of the telomere (Hirai et al., 2012). Finally, chromatin immunoprecipitation experiments in *S. cerevisiae*; *S. pombe* and human cells show that telomerase arrives at telomeres late in S phase. (Bianchi and Shore, 2007b; Hirai et al., 2012; Moser et al., 2009; Smith et al., 2003; Taggart et al., 2002). These experiments are all consistent with the model that telomerase arrives at the telomere with the replication fork in very late S phase. With a new a framework for understanding the linkage of replication and telomere length, previous experimental results can be re-interpreted. For example proteins that were proposed to “recruit” telomerase to the telomere 3’ end ssDNA overhang, such as Cdc13 and Est1, (Evans and Lundblad, 1999) instead may recruit telomerase to the ssDNA created at a telomeric replication fork.

### Lagging strand DNA polymerases and Okazaki fragment processing are linked to telomere elongation

Over 30 years ago, telomere length was shown to be altered by mutations in components of lagging strand DNA synthesis (Carson and Hartwell, 1985). Lagging strand replication occurs by synthesis of short stretches of DNA, called Okazaki fragments, followed by their maturation and ligation to generate a continuous DNA strand (Kurth and O'Donnell, 2013). Each Okazaki fragment begins with synthesis of a primer by the DNA polymerase alpha/primase complex; PCNA is then loaded and recruits DNA polymerase delta (Langston et al., 2009; O'Donnell et al., 2013). During Okazaki fragment maturation, the nascent DNA/RNA strand is processed by Fen1 and Dna2 and joined to the upstream newly synthesized DNA (Balakrishnan and Bambara, 2013). After ligation PCNA must then be unloaded from the newly synthesized DNA (Kubota et al., 2013). Strikingly, mutations in many of the components of lagging strand synthesis affect telomere length.

In *S. cerevisiae* specific hypomorphic alleles of *POL1* (DNA Polymerase alpha) cause
excessive telomere elongation and increased telomeric single stranded DNA (Adams Martin et al., 2000; Carson and Hartwell, 1985). Mutations in genes encoding DNA primase (Pol12), Dna2 and Fen1 that process Okazaki fragments also increase single stranded DNA and telomere length (Budd et al., 2006; Grossi et al., 2004; Parenteau and Wellinger, 1999). Also, mutations in Pif1, a helicase involved in Okazaki fragment maturation (Bochman et al., 2010; Budd et al., 2006), have long telomeres (Schulz and Zakian, 1994). Mutations in the canonical Replication Factor C (RFC), which loads PCNA (Adams and Holm, 1996) as well as in an alternative RFC, composed of ELG1, Ctf18 and Rad24, which unloads PCNA, cause significant telomere elongation (Kanellis et al., 2003; Kubota et al., 2015). This association of lagging strand replication components, including DNA polymerase alpha and RFC, as well as FEN1 to telomere length is conserved across eukaryotes (Dahlen et al., 2003; Derboven et al., 2014; Sampathi et al., 2009; Takashi et al., 2009). The mechanism by which impairment of lagging strand synthesis might lead to telomere *elongation* is not clear, and might seem counterintuitive. Perhaps components of fork stabilization complex stabilize telomerase association with a stalled fork. While the mechanism is not clear, the mechanistic link between lagging strand synthesis and telomere length was further supported by Gottschling’s group. They found that DNA Polymerases alpha and delta and DNA primase are each absolutely required for *de novo* telomere addition by telomerase (Diede and Gottschling, 1999). Examining this link in the context of telomerase association with the fork might shed light on the mechanism of telomere length regulation.

### Telomere specific RPA involved in telomere maintenance

The identification of a telomere specific RPA in yeast further strengthens the link between replication and telomere length. RPA is the eukaryotic single stranded DNA binding protein that binds the single stranded DNA behind the helicase at the fork (Brill and Stillman, 1989; Wold et al., 1989). The RPA complex is a trimer containing RPA70, RPA32 and RPA14 that binds to single stranded DNA and is required for DNA replication (Brill and Stillman, 1991; Erdile et al., 1990). Work from the Lundblad lab first suggested that Cdc13, and its two binding partners Stn1 and Ten1, form a trimeric complex protein resembling RPA (Gao et al., 2007). The similarity to RPA was confirmed by the crystal structure (Gelinas et al., 2009) indicating that Cdc13/Stn1/Ten1 form an alternative, telomere specific, RPA complex, termed t-RPA.

This telomere specific t-RPA (also called CST) is conserved across eukaryotes. In humans, Xenopus and Arabidopsis the large subunit CTC1 does not share sequence identity with *CDC13*, but they do have structural similarities (Miyake et al., 2009; Price et al., 2010; Surovtseva et al., 2009). Stn1 and Ten1 are more conserved and the crystal structure of the human STN1-TEN1 sub complex shows structural conservation with the *S. cerevisiae* t-RPA (Bryan et al., 2013). Mutations in CTC1 cause telomere shortening in patients with human telomere syndromes (Anderson et al., 2012) highlighting its role in mammalian length regulation. siRNA disruption of Stn1 and Ten1 increase telomere length in cultured cells (Bryan et al., 2013). These studies suggest the evolutionally conserved t-RPA (also known as CST) complex plays an essential role in telomere length regulation.

The t-RPA (CST) interacts with polymerase alpha/primase, once again linking lagging strand synthesis with telomere length regulation. In *S. cerevisiae* Cdc13 interacts with the DNA polymerase alpha/primase complex (Nugent et al., 1996; Qi and Zakian, 2000). A co-crystal shows binding of the Cdc13 N-terminal OB fold domain to DNA polymerase alpha (Sun et al., 2011). Stn1 and Ten1 in human cells were first identified as DNA polymerase alpha/primase accessory proteins (Goulian et al., 1990) and biochemical reconstitution shows that the yeast t-RPA (CST) complex can stimulate DNA primase activity *in vitro* (Lue et al., 2014), providing functional evidence for a role in DNA replication as well as telomere length regulation.

### Conserved interaction of RPA and telomerase

If Cdc13 is a part of an alternative RPA, how do we reconcile this with its proposed role in binding of the telomere G strand overhang and providing end protection? The fact that Cdc13 is not needed for end protection outside of S-phase (Vodenicharov et al., 2010) and that ChiP experiments show that Cdc13 binds telomeres almost exclusively in S-phase, calls into question the model that Cdc13 is constitutively bound to the telomeric G-stand overhang. Cdc13 (and the whole t-RPA complex) was proposed to associate with the single stranded telomeric DNA at the replication fork as it passes though the telomere (Gao et al., 2010). Indeed Cdc13 does bind specifically to the lagging strand during telomere replication (Faure et al., 2010). Lundblad has proposed a model in which t-RPA is required to facilitate replication though telomeric DNA and prevent replication fork collapse (CSHL telomere meeting 2013 abstract, and V. Lundblad personal communication). t-RPA binding to DNA polymerase alpha suggests it may also participate directly in telomere lagging strand replication. If Cdc13 associates with single stranded telomeric DNA just behind the fork, it would explain the finding that Cdc13 is only found at telomeres in late S-phase. Telomeres replicate late and so the single stranded telomeric DNA would only be exposed on the lagging strand late in S-phase. The association of Cdc13 with polymerase alpha put this alternative t-RPA squarely at the replication fork.

### Telomerase associates with t-RPA

We have known for over 15 years that CDC13 binds telomerase through interaction with Est1 (Evans and Lundblad, 1999). The recent CryoEM structure of the *Tetrahymena* telomerase holoenzyme also directly links telomerase to t-RPA. The structure shows that two distinct t-RPA complexes are bound with TERT in the telomerase holoenzyme (Jiang et al., 2015). These studies provide compelling evidence that telomerase may associate with the replication fork though its interaction with t-RPA, and thus this complex is an one candidate for a factor that might mediate telomerase association with the replication fork though there could be more than one interaction of telomerase at the fork.

As discussed earlier, in ciliates, telomerase travels with replication bands that represent synchronous replication forks (Fang and Cech, 1995). The telomere binding protein TBPα/TBPβ from *Oxytricha nova,* which was the first such terminal protein identified, (Gottschling and Zakian, 1986; Hicke et al., 1990; Price and Cech, 1989) also travels with the fork in *Oxytricha* (Fang and Cech, 1995) as well as in and *Euplotes* (Skopp et al., 1996). This further suggests that TBPα/TBPβ may be part of a tRPA that associates with telomerase.

## Conclusions

The protein counting model explains some of the experimental data on the negative regulation of telomere length, but it does not account for the role of origin firing, or lagging strand synthesis, in regulating length. When a model is drawn many times in papers and review articles it influences how scientists interpret their experiments. Although many aspects of the replication fork model presented here still need to be tested, it gives the field a different way of thinking about telomere length regulation, and allows for new interpretation of past experiments. Models serve their purpose best when they are tested, shot down, modified or replaced by new models that better fit the experimental evidence. This is how science works best.

## Acknowledgments

I appreciate input and feedback I have had from a number of people over the last year while developing and refining the ideas presented here. First the Greider and Armanios and Holland labs gave feedback at several group meetings and the MBG department faculty provided input at faculty chalk talk. I am indebted to several people in the replication field, Thomas Kelly, James Berger, Steve Bell and Bruce Stillman for talking over some of these issues and reading the manuscript, and to Jeremy Nathans for encouragement to publish this working model. Work in the Greider lab is supported by the NIH grants AG009383 and CA160300 and by Allegheny Health Network Cancer Research Fund.

